# Simulation of Stand-to-Sit Biomechanics for Design of Lower-Limb Exoskeletons and Prostheses with Energy Regeneration

**DOI:** 10.1101/801258

**Authors:** Brock Laschowski, Reza Sharif Razavian, John McPhee

## Abstract

Although regenerative actuators can extend the operating durations of robotic lower-limb exoskeletons and prostheses, these energy-efficient powertrains have been exclusively designed and evaluated for continuous level-ground walking.

**Objective:** Here we analyzed the lower-limb joint mechanical power during stand-to-sit movements using inverse dynamic simulations to estimate the biomechanical energy available for electrical regeneration.

**Methods:** Nine subjects performed 20 sitting and standing movements while lower-limb kinematics and ground reaction forces were measured. Subject-specific body segment parameters were estimated using parameter identification, whereby differences in ground reaction forces and moments between the experimental measurements and inverse dynamic simulations were minimized. Joint mechanical power was calculated from net joint torques and rotational velocities and numerically integrated over time to determine joint biomechanical energy.

**Results:** The hip produced the largest peak negative mechanical power (1.8 ± 0.5 W/kg), followed by the knee (0.8 ± 0.3 W/kg) and ankle (0.2 ± 0.1 W/kg). Negative mechanical work from the hip, knee, and ankle joints per stand-to-sit movement were 0.35 ± 0.06 J/kg, 0.15 ± 0.08 J/kg, and 0.02 ± 0.01 J/kg, respectively.

**Conclusion and Significance:** Assuming an 80-kg person and previously published regenerative actuator efficiencies (i.e., maximum 63%), robotic lower-limb exoskeletons and prostheses could theoretically regenerate ~26 Joules of total electrical energy while sitting down, compared to ~19 Joules per walking stride. Given that these regeneration performance calculations are based on healthy young adults, future research should include seniors and/or rehabilitation patients to better estimate the biomechanical energy available for electrical regeneration among individuals with mobility impairments.

## I. Introduction

OVER 12 million people in the United States alone have mobility impairments resulting from stroke, spinal cord injury, and other neuromusculoskeletal diseases [1]. There are approximately 2 million Americans with limb amputations [2]; these numbers are expected to increase with the emergent aging population and growing incidences of cancer and diabetes [1]–[3]. Robotic lower-limb exoskeletons and prostheses can help seniors and rehabilitation patients perform movements that involve net positive mechanical work (e.g., sit-to-stand) by mimicking their amputated or unimpaired biological muscles [4]–[9]. However, these biomechatronic devices have typically required significant electrical power and heavy onboard batteries to facilitate daily functioning [5]–[6], [10]–[11]. For instance, robotic knee prostheses under research and development have consumed 43 ± 30 W of electricity during level-ground walking; provided only 3.1 ± 2.2 hours of maximum operation; and weighed 4.0 ± 1.1 kg [4], [6]. Robotic lower-limb exoskeletons have provided only 1-5 hours of maximum operation [1]. Portable electricity has been considered a leading challenge to developing robotic exoskeletons for real-world environments [1], [10]–[11]. Research into energy-efficient biomechatronic design and control systems is thus warranted.

Electrical energy regeneration is a potential solution to the aforementioned shortcomings. Human joints can produce both negative mechanical power (braking) and positive mechanical power (motoring) [12]. During level-ground walking, the human knee joint resembles a damper mechanism, performing net negative mechanical work via energy dissipation, and the ankle joint resembles an actuating motor, performing net positive mechanical work and generating forward propulsion [5], [10]–[12]. These characteristic human walking biomechanics are illustrated in Fig. 1. Similar to electric and hybrid electric vehicles [13]–[14], several biomechatronic knee designs have incorporated regenerative actuators that convert otherwise dissipated joint biomechanical energy during negative work movements into electrical energy by reversing the direction of operation [5], [15]–[30]. Such bidirectional power flow during motoring and generating operations requires backdriveable actuator-transmission systems with low mechanical impedance [31]–[36]. These energy-efficient powertrain designs can enable lighter onboard batteries and/or extend the operating durations between recharging. For socket-suspended lower-limb prostheses, decreasing the onboard battery weight can also minimize 1) the metabolic power consumption during walking, and 2) discomfort from excessive tugging on the human-socket interface [2].

**Fig. 1.**
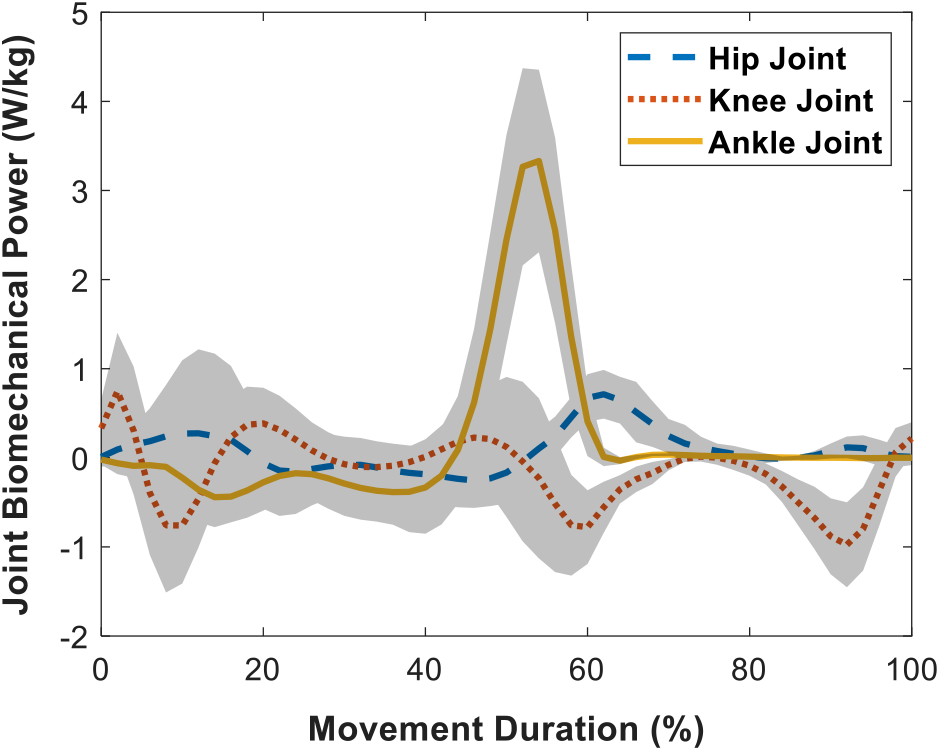
Hip, knee, and ankle joint mechanical power per level-ground walking stride normalized to total body mass. Data were taken from [12]. The uncertainties are ± one standard deviation across different subjects (n=19). The movement trajectories begin and end with heel-strike.

Previous studies of lower-limb exoskeletons and prostheses with regenerative actuators have focused exclusively on continuous level-ground walking [15]–[30], [33], [37]–[39]. However, seniors and rehabilitation patients typically exhibit slower walking speeds (e.g., ~24% reduction from 25 to 75 years) and take fewer steps/day (e.g., ~75% reduction from 60 to 85 years) [3], therein limiting the potential for electrical regeneration from level-ground walking. Conversely, sitting and standing movements can be considered more applicable activities of individuals with mobility impairments. Healthy young adults perform ~60 sitting and standing movements per day [40]. Several lower-limb exoskeletons and prostheses have been designed and evaluated for sitting and standing movements (although without regenerative actuators) [7]–[9], [40]–[46]. Regenerating energy while sitting down represents an unexplored and potentially viable method to supplementing that from level-ground walking. Motivated to find new opportunities for energy savings, we analyzed the lower-limb joint mechanical power during stand-to-sit movements using inverse dynamics to estimate the biomechanical energy available for electrical regeneration.

## II. Methods

### A. Motion Capture Experiments

Nine subjects were recruited and provided informed written consent (height: 180 ± 4 cm, body mass: 78 ± 7 kg, age: 25 ± 3 years, sex: male). Each subject performed 20 sitting and standing movements while lower-limb kinematics and ground reaction forces were measured using motion capture cameras and force plates, respectively (see Fig. 2). The seat height was 46 cm. The motion capture cameras (Optotrak, Northern Digital Incorporation, Canada) provided 3D measurements of active marker positions in global coordinates. Active marker systems are generally considered the gold standard in human movement biomechanics [48]. The motion capture cameras and force plates were sampled at 100 Hz and 300 Hz, respectively. For tracking individual body segment positions in the sagittal plane, virtual markers were digitized overlying palpable anatomical landmarks on the right lower-limb, including the lateral malleolus, lateral femoral and tibial condyles, and greater trochanter. These marker positions correspond with those recommended by the International Society of Biomechanics [48]. This study was approved by the University of Waterloo research ethics office.

**Fig. 2.**
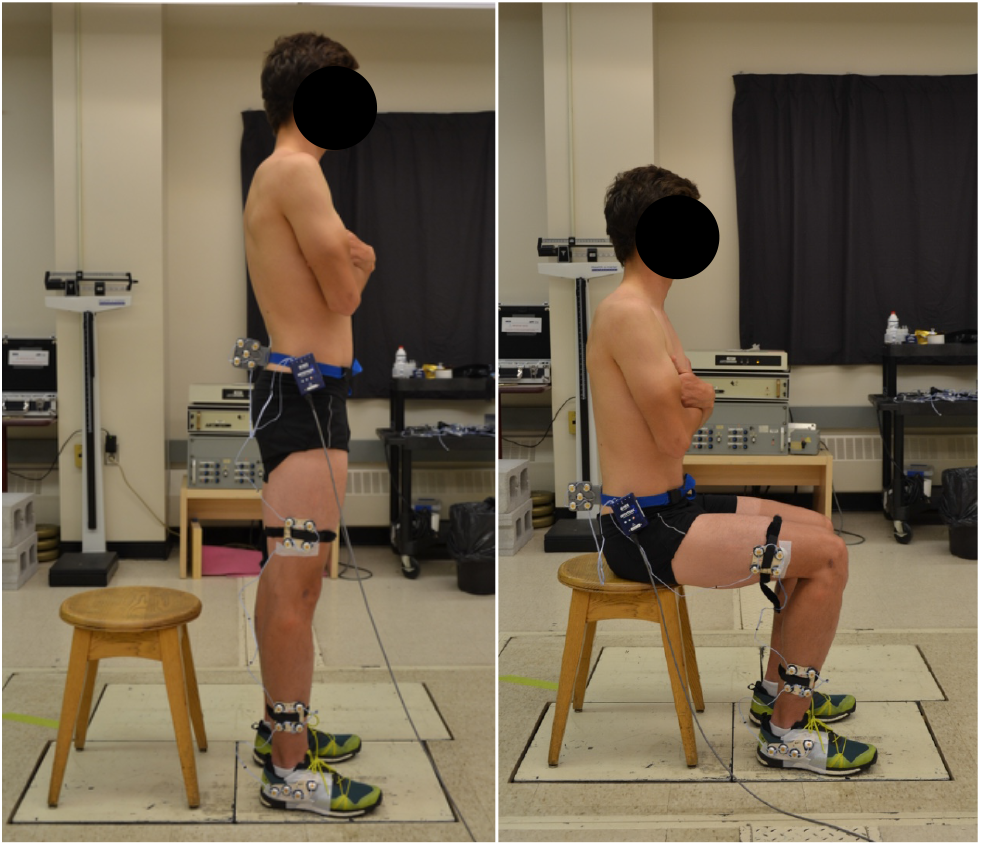
Photographs of the experimental measurements of stand-to-sit movement biomechanics with motion capture cameras and force plates.

### B. Data Processing

Missing marker data were estimated using cubic spline interpolations. The ankle and hip joint centers were assumed at the lateral malleolus and greater trochanter markers, respectively. The estimated knee joint center was the midpoint between the lateral femoral and tibial condyle markers [48]. Piecewise cubic Hermite interpolating polynomials were used to resample and time-normalize the kinematic measurements. Mean line vectors between the ankle and knee joint centers, and knee and hip joint centers, defined the shank and thigh body segment lengths, respectively. Inverse kinematics converted the marker positions to joint coordinates through vector algebra. The ankle angle was the angle between the shank and horizontal axis (Fig. 3). The relative angle between the shank and thigh segments defined the knee angle. Given the relative rotations between the pelvis and HAT segment, the measured pelvis marker-cluster rotations differed from HAT segment rotations. Therefore, the HAT segment was assumed vertical when standing (initial posture) and seated (final posture) and the pelvis angle rotations were assumed to progress linearly throughout the movement. The angle between the thigh and HAT segments defined the hip angle. Joint angles were filtered using 10th-order low-pass Butterworth filters with 5 Hz cut-off frequencies and zero-phase digital filtering [12]. Joint rotational speeds and accelerations were calculated by numerically differentiating the joint angles.

**Fig. 3.**
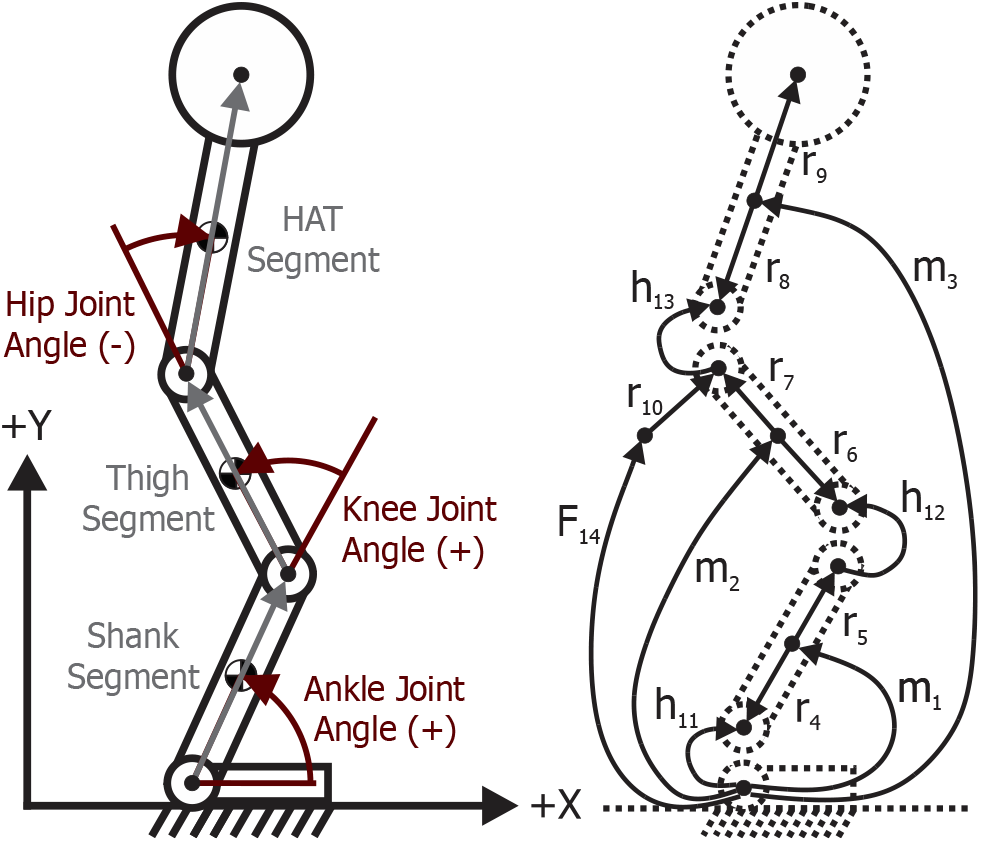
(left) 2D human biomechanical model including hip, knee, and ankle joints and HAT, thigh, and shank segments. (right) Linear graph of the biomechanical model. Edges m_1-3_ are body segment inertias; elements r_4-9_ are constant body-fixed position vectors; edge r_10_ represents the “seat_offset_” position vector fixed to the thigh segment; h_11-13_ are revolute joints; and F_14_ represents the external seat force.

Similar to previous research [41], the pelvis translational velocities, which were estimated from the marker-cluster shown in Fig. 2, were used to segment the sitting and standing movements. These kinematic measurements were filtered using 10th-order low-pass Butterworth filters and 3 Hz cut-off frequencies, and zero-phase digital filtering and moving average smoothing filtering. The sitting and standing movements were segmented when the pelvis translational velocities exceeded a percentage of their maximum values, which were estimated through trial-and-error simulations. Force plate measurements were filtered using 10th-order low-pass Butterworth filters and 30 Hz cut-off frequencies, and zero-phase digital filtering. Piecewise cubic Hermite interpolating polynomials were used to time-normalize the force plate measurements.

### C. Biomechanical Model Design

The human biomechanical system was dynamically modelled using MapleSim (Maplesoft, Canada). The biomechanical model comprised a 2D sagittal-plane inverted triple-pendulum with shank, thigh, and HAT rigid body segments (see Fig. 3). The foot segment was fixed to the ground. The ankle, knee, and hip were modelled as revolute joints. Biological passive joint torques, including stiffness and damping, were ignored since we assumed ideal joints for modelling exoskeleton and prosthetic systems. The biomechanical model had three degrees-of-freedom and was represented by three generalized coordinates with zero algebraic constraints. Given that the foot was fixed to the ground and had relatively small mass, the ground reaction forces corresponded with the ankle joint reaction forces and the ground reaction moments were offset by ankle position relative to the center of pressure. The measured ground reaction forces underneath the seat were applied to the model buttocks when seated. MapleSim automatically generated the multibody system equations symbolically using linear graph theory, therein enabling computationally-efficient dynamic simulations. Fig. 3 also shows the linear graph of the biomechanical model.

### D. Simulation and Parameter Identification

The biomechanical model was driven using the experimental joint kinematics and seat forces. The ankle, knee, and hip joint torques (τ), and ground reaction forces and moment underneath the foot, were calculated from inverse dynamics; this method can be advantageous over bottom-up inverse dynamics because it yields dynamically consistent simulations with zero residual forces/moments on the final body segment (HAT). Human body segment parameters can be estimated using medical imaging [49]–[50], system parameter identification, and/or anthropometric proportionalities from cadaver research [51]. We decided to use parameter identification for better dynamical consistency. The identification involved constrained nonlinear programming (Fmincon, MATLAB) and an interior-point algorithm to estimate the body segment inertial parameters (i.e., mass, center of mass, and moment of inertia). The optimization searched for the parameters that minimized the sum of squared differences in ground reaction forces (GRF) and moments (GRM) between the experimental measurements (m) and inverse dynamic simulations (s) across each time step (i). The optimization multiobjective cost function was:

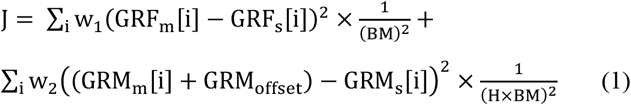

where the two-dimensional GRF vector included both the horizontal (GRF_x_) and vertical (GRF_y_) components, GRM was around the z-axis, BM was body mass, coefficient H = 1 meter, and GRM_offset_ compensated for the distance between the ankle and foot center of pressure according to {GRM_offset_ = GRF_y_ × COP_x_ - GRF_x_ × COP_y_} with COP_x_ and COP_y_ being approximate constant positions of the foot center of pressure relative to the ankle joint. The optimization variables were the shank, thigh, and HAT segment mass, center of mass, and moment of inertia. Other variables included “seat_offset_” and the COP_x_. Seat_offset_ was the distance between the model buttocks (i.e., vertical seat force point-of-application) and hip joint center. COP_y_ is known from the ankle marker height. The optimization was constrained by setting 1) lower and upper bounds on individual variables, and 2) the summed segment masses equaled to the measured total mass. Initial guesses were taken from human anthropometrics and/or were midpoints between upper and lower bounds. Each term in the optimization had equal weights. Stopping criteria for the step size and objective function changes were both 1e-8 between iterations. The experimental and computational methods are summarized in Fig. 4. Once the optimal parameters were found, joint mechanical powers were calculated from joint torques and rotational speeds 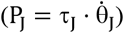 and numerically integrated over time to estimate joint biomechanical energies.

**Fig. 4.**
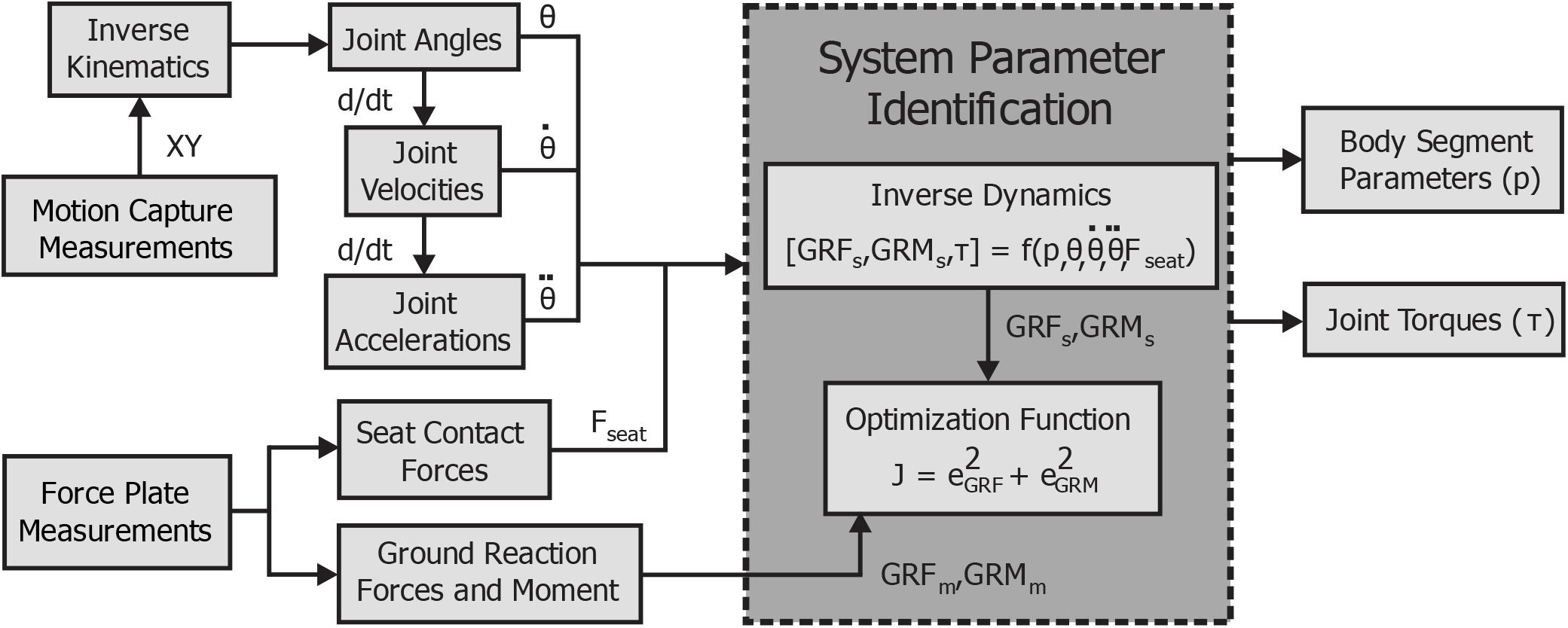
Flow diagram of the experimental and computational methods, including the biomechanical measurements, inverse kinematic and dynamic analyses, and system parameter identification. Nomenclature are defined in the text.

## III. Results

For both kinematic and dynamic results, the movement trajectories were time-normalized to facilitate between and within subject averaging. Fig. 5 shows the calculated hip, knee, and ankle joint angles during stand-to-sit movements from inverse kinematics. Decreasing joint angles represented hip flexion, knee extension, and ankle dorsiflexion, while increasing joint angles represented hip extension, knee flexion, and ankle plantar flexion. The uncertainties are ± one standard deviation across each subject (n=9) and trial (20 trials/subject), totaling 180 individual trials. There were minor variations in the joint kinematics between and within subjects, as demonstrated by the small standard deviations.

**Fig. 5.**
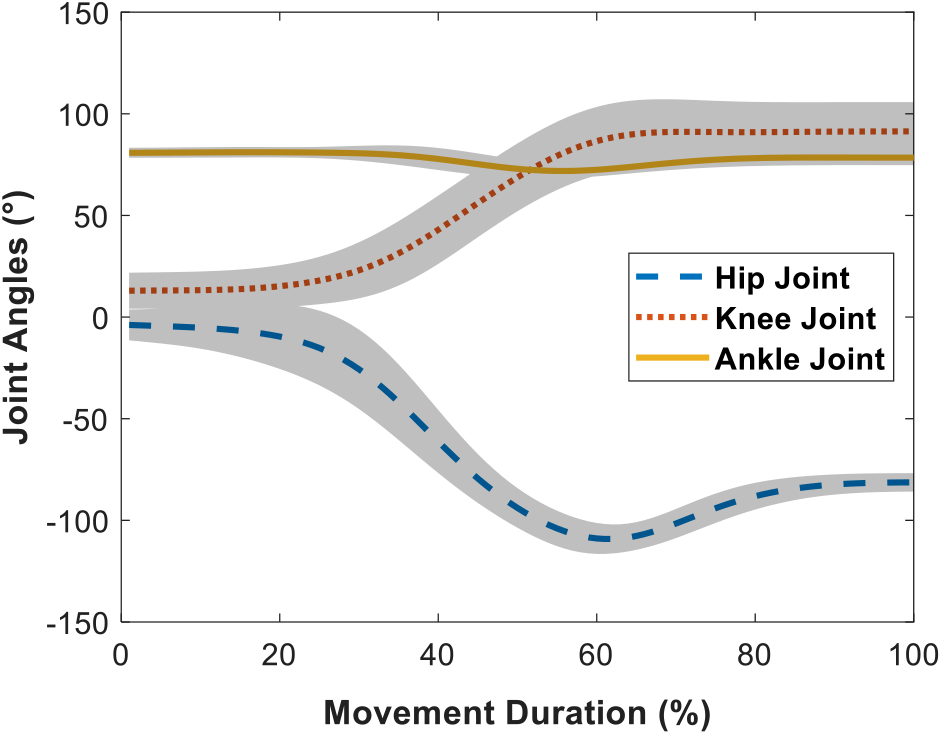
Hip, knee, and ankle joint angles during stand-to-sit movements from inverse kinematics. The uncertainties are ± one standard deviation across each subject (n=9) and trial (20 trials/subject).

Fig. 6 shows the calculated hip, knee, and ankle joint torques from inverse dynamics; the corresponding maximum values were 0.7 ± 0.1 Nm/kg, 1.1 ± 0.3 Nm/kg, and 0.4 ± 0.1 Nm/kg. The calculated hip, knee, and ankle joint mechanical powers during stand-to-sit movements are shown in Fig. 7. The hip produced the largest peak negative mechanical power (1.8 ± 0.5 W/kg), followed by the knee joint (0.8 ± 0.3 W/kg) and ankle joint (0.2 ± 0.1 W/kg). Negative mechanical work from the hip, knee, and ankle joints were 0.35 ± 0.06 J/kg, 0.15 ± 0.08 J/kg, and 0.02 ± 0.01 J/kg, respectively. Subjective feedback from participants indicated that performing stand-to-sit movements was significantly more challenging than sit-to-stand movements, particularly from a balance control perspective. The experimental and simulated biomechanical data were uploaded to IEEE DataPort and are available for download at https://ieee-dataport.org/documents/measurement-and-simulation-human-sitting-and-standing-movement-biomechanics.

**Fig. 6.**
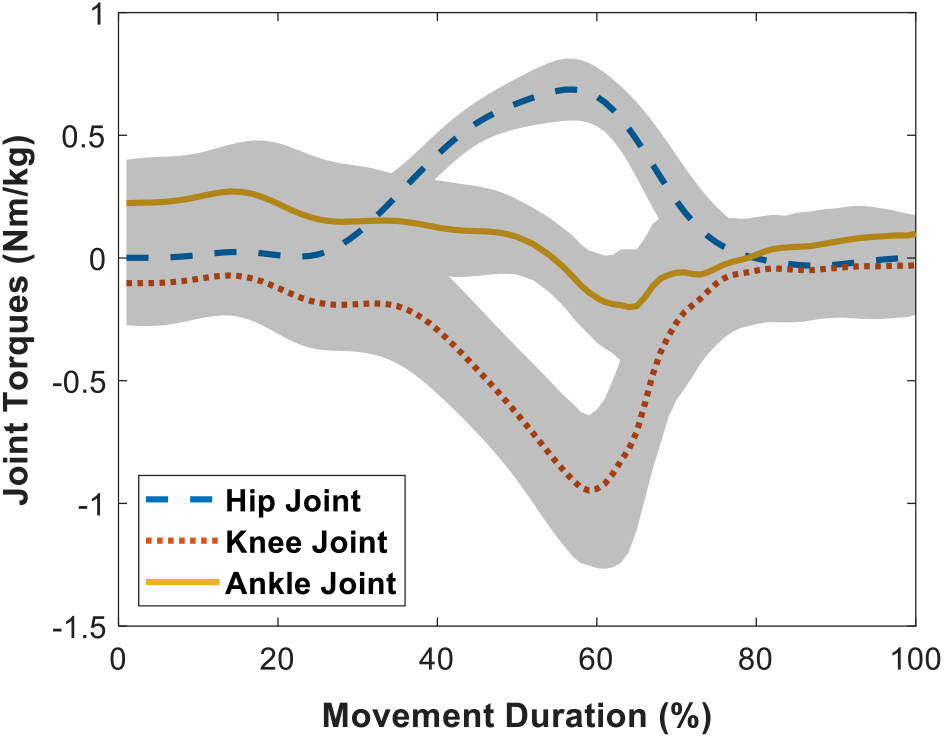
Hip, knee, and ankle joint torques during stand-to-sit movements from inverse dynamics normalized to total body mass. The uncertainties are ± one standard deviation across each subject (n=9) and trial (20 trials/subject).

**Fig. 7.**
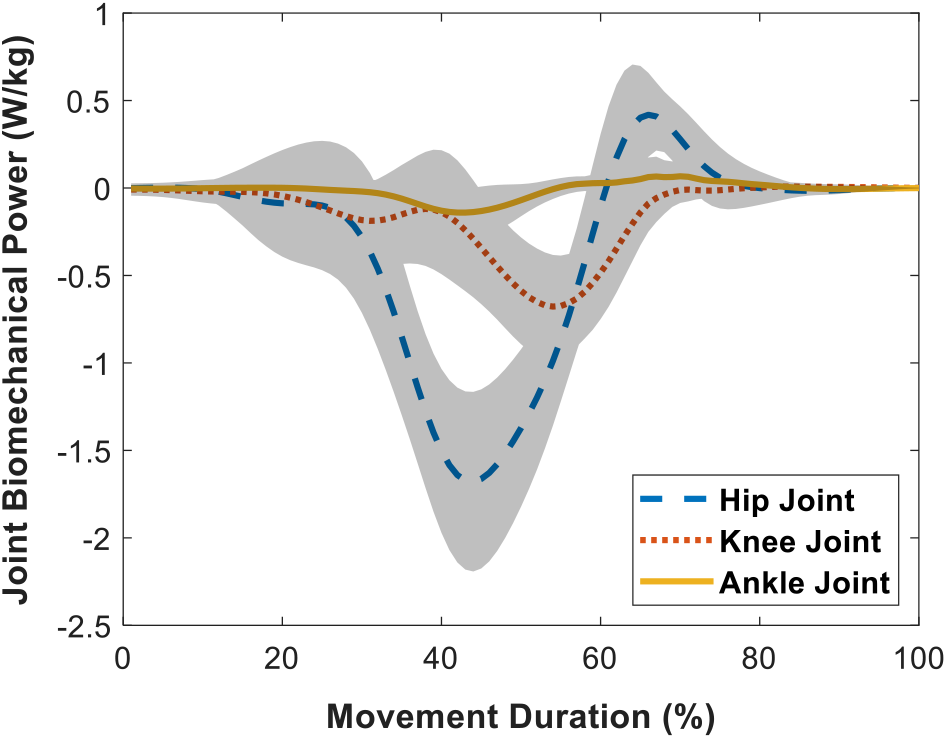
Hip, knee, and ankle joint mechanical power during stand-to-sit movements normalized to total body mass. The uncertainties are ± one standard deviation across each subject (n=9) and trial (20 trials/subject).

## IV. Discussion

Comparable to electric and hybrid electric vehicles [13]–[14], regenerative actuators can extend the operating durations of robotic lower-limb exoskeletons and prostheses by converting the otherwise dissipated joint biomechanical energy during negative work movements into electrical energy for recharging the onboard batteries, hence the term regenerative braking. However, for biomechatronic applications, these energy-efficient powertrains have been exclusively designed and evaluated for level-ground walking [15]–[30], [33], [37]–[39]. Building on previous research, we analyzed the lower-limb joint mechanical power during stand-to-sit movements using inverse dynamic simulations to estimate the biomechanical energy available for electrical regeneration during activities considered more applicable to aging and rehabilitation populations (considering that some individuals perform no locomotor activities other than sitting and standing movements for wheelchair transfers) [3]. The calculated peak negative mechanical powers from the hip, knee, and ankle joints were 1.8 ± 0.5 W/kg, 0.8 ± 0.3 W/kg, and 0.2 ± 0.1 W/kg, respectively. In comparison, experimental measurements with lower-limb prostheses reported 0.7-0.8 W/kg of peak robotic knee joint mechanical power during sitting and standing movements [40], [45]. The strong quantitative agreement between the simulated (0.8 ± 0.3 W/kg) and experimental (0.7-0.8 W/kg) [40], [45] knee joint mechanical powers supported the model validation. The model validation was further corroborated by relatively good agreements in maximum knee joint torques between our biomechanical simulations (1.1 ± 0.3 Nm/kg) and previous research on robotic lower-limb exoskeletons and prostheses during sitting and standing movements (0.8-1.0 Nm/kg) [40],[42]–[44]. Note that these joint torques are high enough to dynamically backdrive an actuator-transmission system (i.e., requiring 1-3 Nm torque) [32]–[34], [36] and thus capable of regenerating electrical energy while sitting down.

The hip joint performed the most negative mechanical work during stand-to-sit movements (0.35 ± 0.06 J/kg), followed by the knee (0.15 ± 0.08 J/kg) and ankle (0.02 ± 0.01 J/kg). Although the hip performs the most negative mechanical work, and therefore has the greatest potential for electrical regeneration, most lower-limb exoskeletons and prostheses with regenerative actuators have featured knee-centered designs [15]–[20], [22]–[30], [33], [37]–[39]. Fig. 8 presents an example powertrain system model used to estimate regeneration performance. A standard electric motor converts electrical power (VI) to mechanical power 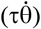. When backdriven, the motor works like a generator, converting mechanical power to electrical power. Regenerative actuator efficiency (η) is defined as the percentage of mechanical-to-electrical power conversion. Assuming an 80-kg person and previously published regenerative actuator efficiencies (η = maximum 63%) [19], [25]–[26], [52]–[53], robotic lower-limb exoskeletons and prostheses could theoretically regenerate ~26 J of total electrical energy while sitting down. Backdriving the same regenerative actuator model using Winter’s walking data [12], ~19 J of total electrical energy could be regenerated per stride. These calculations assume 1) bidirectionally symmetric and constant (i.e., torque and velocity independent) actuator efficiencies, 2) losses only from the actuator-transmission system (i.e., Joule heating and friction), and 3) electrical regeneration over the entire negative joint mechanical power range. Tables 1 and 2 summarize the individual joint biomechanical energies during level-ground walking and stand-to-sit movements, respectively.

**Fig. 8.**
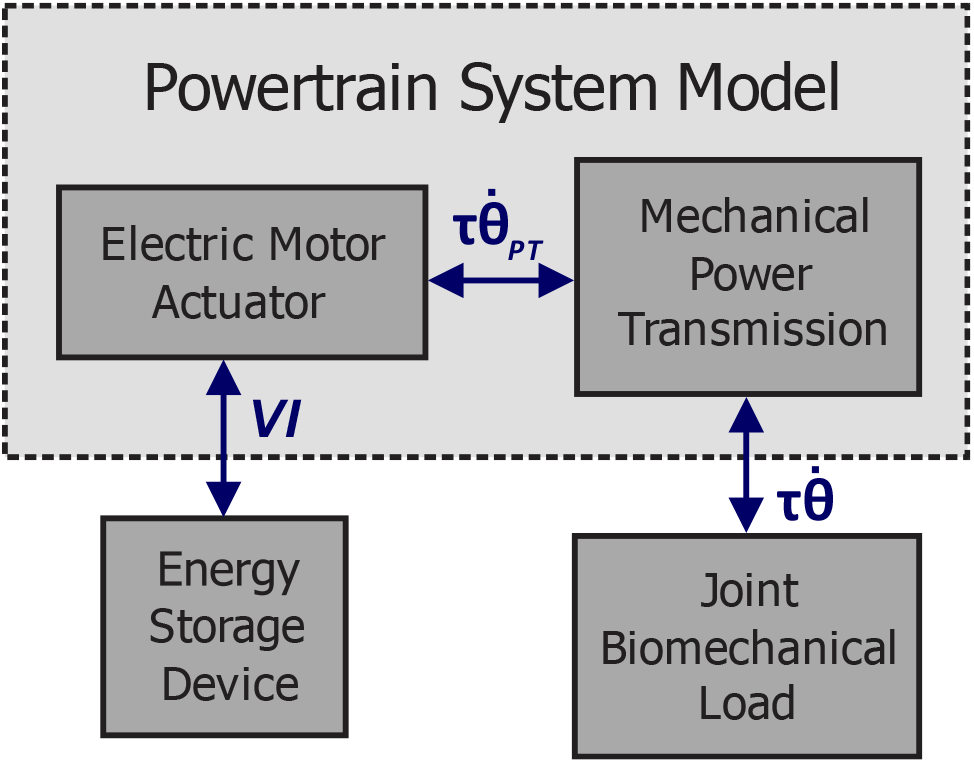
Example of an exoskeleton/prosthesis powertrain system model, including an energy storage device, electric motor actuator, mechanical power transmission, and biomechanical load. The arrows represent the bidirectional flow of electrical (VI) and mechanical 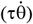 power during motoring and generating operations.

**Table 1.** Hip, knee, and ankle joint mechanical work per level-ground walking stride [12]. The results are averages across multiple subjects (n=19) and normalized to total body mass (J/kg). Total Work is the combined biomechanical energies from each joint and Net Joint Work is the net lower-limb joint mechanical work performed on the system.

|  | Positive Work | Negative Work | Net Joint Work |
| --- | --- | --- | --- |
| Hip Joint | 0.130 | -0.049 | 0.081 |
| Knee Joint | 0.074 | -0.223 | -0.149 |
| Ankle Joint | 0.313 | -0.110 | 0.203 |
| Total Work | 0.517 | -0.382 | 0.135 |

**Table 2.** Hip, knee, and ankle joint mechanical work per stand-to-sit movement. The results are averages across multiple subjects (n=9) and trials (20 trials/subject) and normalized to total body mass (J/kg). Total Work is the combined biomechanical energies from each joint and Net Joint Work is the net lower-limb joint mechanical work performed on the system.

|  | Positive Work | Negative Work | Net Joint Work |
| --- | --- | --- | --- |
| Hip Joint | 0.038 | -0.347 | -0.309 |
| Knee Joint | 0.001 | -0.147 | -0.146 |
| Ankle Joint | 0.011 | -0.022 | -0.011 |
| Total Work | 0.050 | -0.516 | -0.466 |

Integrating the positive mechanical powers in Fig. 1 provides insight into the energetic requirements (battery consumption) of robotic lower-limb exoskeletons and prostheses during human locomotion. Based on these calculations, level-ground walking requires ~0.52 J/kg of total positive lower-limb joint mechanical work per stride to generate forward propulsion, which equates to ~66 J of electrical energy, assuming the aforementioned body mass and powertrain system model. Using a single rechargeable lithium-ion battery (2.6 Ah and 24 V; total battery capacity of 224,640 J) [21], lower-limb exoskeletons and pros-theses could theoretically walk ~3,404 steps between recharging. Regenerating electrical energy during level-ground walking and stand-to-sit movements (i.e., assuming 60 movements per day) [40] could therefore individually extend the operating durations by an additional 40% (~4,780 total steps) and 0.7% (~3,428 total steps). In other words, level-ground walking and stand-to-sit motions could regenerate ~64,676 J and ~1,560 J of total electrical energy each day, respectively. Here we assume sufficient power electronics to control the bidirectional flow of electrical power between the motor and onboard battery. Although regeneration during level-ground walking can regenerate more electrical energy than stand-to-sit movements over an extended period (i.e., assuming healthy biomechanical data), there are projected advantages to energy regeneration from stand-to-sit movements, as subsequently discussed.

Control of regenerative actuators is notoriously challenging [5]. Regeneration control during stand-to-sit movements would theoretically have higher tolerances to reference tracking errors since the lower-limb joint biomechanical energies are almost entirely negative (see Fig. 7). In comparison, energy regeneration during level-ground walking requires more robust controls since the lower-limb joint biomechanical energies are intermittent, multidirectional, and time-varying (see Fig. 1). Inaccurate and/or delayed reference tracking could result in regenerating electrical energy during periods of positive mechanical work. Unlike regenerative braking, energy regeneration during positive mechanical work would require human muscles to actively backdrive the actuator-transmission system and perform more positive mechanical work, therein increasing metabolic power consumption and decreasing overall efficiency [19]–[20], [25]–[26]. These energetic consequences are especially pertinent to seniors and rehabilitation patients who already exhibit more inefficient walking [3]. Ultimately, there are advantages and disadvantages to regeneration from different human movements (e.g., stand-to-sit movements may regenerate less electrical energy per day than level-ground walking, but facilitate more robust reference tracking and control). For maximum efficiency and battery performance, robotic lower-limb exoskeletons and prostheses should regenerate energy from many different negative work movements (e.g., walking, sitting down, and ramp and stair descent).

We want to acknowledge that our regeneration performance calculations are based entirely on healthy young adults, therein requiring numerous assumptions and (potentially questionable) extrapolations to aging and rehabilitation populations. Healthy adults exhibit self-selected walking speeds around 1.35 m/s and walk between 6,000 and 13,000 steps/day [3]. In contrast, seniors typically walk slower and shorter distances. Approximately 50% of individuals over 65 years walk less than 5000 steps/day [3]. These movement patterns are further worsened in patients with neuromusculoskeletal diseases. For instance, patients with incomplete spinal cord injuries walk ~1,640 steps/day [3]. Such differences relative to healthy young adults have implications on regenerative actuator performance.

Empirical studies of lower-limb exoskeletons and prostheses with regenerative actuators have shown a positive correlation between walking speed and both electrical energy regeneration and efficiency (i.e., faster walking generates more electricity and more efficiently) [15]–[16], [21], [23], [25]–[26], [33]. For an assumed back EMF constant, an electric motor will generate a voltage proportional to the rotational speed. Slower walking speeds, such as that characteristic of seniors and rehabilitation patients [3], would backdrive the motor with lower rotational speeds and therefore generate less electrical power. Motors are also typically less efficient when generating torques at lower speeds because of Joule heating. For instance, a recent study [33] showed that increasing walking speed from 0.9 m/s to 1.6 m/s increased power conversion efficiency from 40% to 59%. Taking into account the differences in movement biomechanics between healthy adults and individuals with mobility impairments, and their implications on electrical energy regeneration and efficiency, future research should include seniors and/or rehabilitation patients to improve our regeneration performance estimates.

## V. Conclusion

Regenerative actuators can increase the energy-efficiency and extend the operating durations of robotic exoskeletons and prostheses by converting otherwise dissipated biomechanical energy during negative work movements into electrical energy for battery recharging. However, previous research has focused exclusively on continuous level-ground walking. Motivated to expand the state-of-the-science and find new opportunities for energy savings, we analyzed the lower-limb joint mechanical power during stand-to-sit movements using inverse dynamic simulations to estimate the biomechanical energy available for electrical regeneration. Our simulations showed that the hip joint produced the largest peak negative mechanical power (1.8 ± 0.5 W/kg), followed by the knee (0.8 ± 0.3 W/kg) and ankle (0.2 ± 0.1 W/kg). Negative mechanical work from the hip, knee, and ankle joints per stand-to-sit movement were 0.35 ± 0.06 J/kg, 0.15 ± 0.08 J/kg, and 0.02 ± 0.01 J/kg, respectively. Back-driving a regenerative actuator system on each lower-limb joint using these biomechanical inputs, robotic exoskeletons and prostheses could theoretically regenerate ~26 Joules of total electrical energy while sitting down, compared to ~19 Joules per walking stride. However, these regeneration performance calculations are based on healthy young adults. To more accurately estimate the biomechanical energy available for electrical regeneration with lower-limb exoskeletons and prostheses, in addition to the operational performances of the regenerator actuator and onboard batteries, future research should expand our analyses to include seniors and/or rehabilitation patients.

## Acknowledgments

This research was funded by the Natural Sciences and Engineering Research Council of Canada (NSERC), the University of Waterloo Engineering Excellence PhD Fellowship, and Dr. John McPhee’s Canada Research Chair in Biomechatronic System Dynamics. We wish to thank Dr. Stacey Acker (University of Waterloo, Canada) and Mr. Mark Charlet (Université Laval, Canada) for assisting with the biomechanical measurements.

## References

[1] A. J. Young and D. P. Ferris, “State of the art and future directions for lower limb robotic exoskeletons,” IEEE Transactions on Neural Systems and Rehabilitation Engineering, vol. 25, no. 2, pp. 171–182, 2017, DOI: 10.1109/TNSRE.2016.2521160.

[2] A. Maryniak, B. Laschowski, and J. Andrysek, “Technical overview of osseointegrated transfemoral prostheses: Orthopedic surgery and implant design centered,” ASME Journal of Engineering and Science in Medical Diagnostics and Therapy, vol. 1, no. 2, pp. 020801–020801-7, 2018, DOI: 10.1115/1.4039105.

[3] M. Grimmer, R. Riener, C. J. Walsh, and A. Seyfarth, “Mobility related physical and functional losses due to aging and disease - A motivation for lower limb exoskeletons,” Journal of NeuroEngineering and Rehabilitation, vol. 16, 2019, DOI: 10.1186/s12984-018-0458-8.

[4] B. Laschowski and J. Andrysek, “Electromechanical design of robotic transfemoral prostheses,” in Proceedings of the ASME International Design Engineering Technical Conferences and Computers and Information in Engineering Conference, Quebec City, Canada, 2018, pp. V05AT07A054, DOI: 10.1115/DETC2018-85234.

[5] B. Laschowski, J. McPhee, and J. Andrysek, “Lower-limb prostheses and exoskeletons with energy regeneration: Mechatronic design and optimization review,” ASME Journal of Mechanisms and Robotics, vol. 11, no. 4, pp. 040801–040801-8, 2019, DOI: 10.1115/1.4043460.

[6] D. S. Pieringer, M. Grimmer, M. F. Russold, and R. Riener, “Review of the actuators of active knee prostheses and their target design outputs for activities of daily living,” in Proceedings of the IEEE International Conference on Rehabilitation Robotics (ICORR), London, UK, July 17-20, 2017, pp. 1246–1253, DOI: 10.1109/ICORR.2017.8009420.

[7] A. M. Simon, N. P. Fey, K. A. Ingraham, A. J. Young, and L. J. Hargrove, “Powered prosthesis control during walking, sitting, standing, and non-weight bearing activities using neural and mechanical inputs,” in Proceedings of the International IEEE/EMBS Conference on Neural Engineering (NER), San Diego, USA, November 6-8, 2013, pp. 1174–1177, DOI: 10.1109/NER.2013.6696148.

[8] J. Skelton, S. K. Wu, and X. Shen, “Design of a powered lower-extremity orthosis for sit-to-stand and ambulation assistance,” ASME Journal of Medical Devices, vol. 7, no. 3, pp. 030910–030910-2, 2013, DOI: 10.1115/1.4024489.

[9] S. Thapa, H. Zheng, G. F. Kogler, and X. Shen, “A robotic knee orthosis for sit-to-stand assistance,” in Proceedings of the ASME Dynamic Systems and Control Conference (DSCC), Minneapolis, USA, 2016, pp. V001T07A004, DOI: 10.1115/DSCC2016-9891.

[10] A. M. Dollar and H. Herr, “Active orthoses for the lower-limbs: Challenges and state of the art,” in Proceedings of the IEEE International Conference on Rehabilitation Robotics (ICORR), Noordwijk, Netherlands, June 13-15, 2017, pp. 968–977, DOI: 10.1109/ICORR.2007.4428541.

[11] A. M. Dollar and H. Herr, “Lower extremity exoskeletons and active orthoses: Challenges and state-of-the-art,” IEEE Transactions on Robotics, vol. 24, no. 1, pp. 144–158, 2008, DOI: 10.1109/TRO.2008.915453.

[12] D. A. Winter, “The Biomechanics and Motor Control of Human Gait: Normal, Elderly, and Pathological,” Waterloo Biomechanics, Canada, 1998.

[13] R. Razavian, N. L. Azad, and J. McPhee, “On real-time optimal control of a series hybrid electric vehicle with an ultra-capacitor,” in Proceedings of the American Control Conference (ACC), Montreal, Canada, June 27-29, 2012, pp. 547–552, DOI: 10.1109/ACC.2012.6314831.

[14] G. Rizzoni and H. Peng, “Hybrid and electrified vehicles: The role of dynamics and control,” ASME Mechanical Engineering Magazine, vol. 135, no. 03, pp. S10–S17, 2012, DOI: 10.1115/1.2013-MAR-5.

[15] J. Andrysek and G. Chau, “An electromechanical swing-phase-controlled prosthetic knee joint for conversion of physiological energy to electrical energy: Feasibility study,” IEEE Transactions on Biomedical Engineering, vol. 54, no. 12, pp. 2276–2283, 2007, DOI: 10.1109/TBME.2007.908309.

[16] J. Andrysek, T. Liang, and B. Steinnagel, “Evaluation of a prosthetic swing-phase controller with electrical power generation,” IEEE Transactions on Neural Systems and Rehabilitation Engineering, vol. 17, no. 4, pp. 390–396, 2009, DOI: 10.1109/TNSRE.2009.2023292.

[17] T. Barto and D. Simon, “Neural network control of an optimized regenerative motor drive for a lower-limb prosthesis,” in Proceedings of the American Control Conference (ACC), Seattle, USA, May 24-26, 2017, pp. 5330–5335, DOI: 10.23919/ACC.2017.7963783.

[18] E. Bolivar, S. Rezazadeh, and R. Gregg, “A general framework for minimizing energy consumption of series elastic actuators with regeneration,” in Proceedings of the ASME Dynamic Systems and Control Conference (DSCC), Tysons, USA, 2017, pp. V001T36A005, DOI: 10.1115/DSCC2017-5373.

[19] J. M. Donelan, Q. Li, V. Naing, J. A. Hoffer, D. J. Weber, and A. D. Kuo, “Biomechanical energy harvesting: Generating electricity during walking with minimal user effort,” Science, vol. 319, no. 5864, pp. 807–810, 2008, DOI: 10.1126/science.1149860.

[20] J. M. Donelan, V. Naing, and Q. Li, “Biomechanical energy harvesting,” in Proceedings of the IEEE Radio and Wireless Symposium, San Diego, USA, January 18-22, 2009, pp. 1–4, DOI: 10.1109/RWS.2009.4957269.

[21] Y. Feng, J. Mai, S. K. Agrawal, and Q. Wang, “Energy regeneration from electromagnetic induction by human dynamics for lower extremity robotic prostheses,” IEEE Transactions on Robotics, 2020, DOI: 10.1109/TRO.2020.2991969.

[22] E. Gualter Dos Santos and H. Richter, “Modeling and control of a novel variable-stiffness regenerative actuator,” in Proceedings of the ASME Dynamic Systems and Control Conference (DSCC), Atlanta, USA, 2018, pp. V002T24A003, DOI: 10.1115/DSCC2018-9054.

[23] P. Khalaf, H. Warner, E. Hardin, H. Richter, and D. Simon, “Development and experimental validation of an energy regenerative prosthetic knee controller and prototype,” in Proceedings of the ASME Dynamic Systems and Control Conference (DSCC), Atlanta, USA, 2018, pp. V001T07A008, DOI: 10.1115/DSCC2018-9091.

[24] B. H. Kim and H. Richter, “Energy regeneration-based hybrid control for transfemoral prosthetic legs using four-bar mechanism,” in Proceedings of the Annual Conference of the IEEE Industrial Electronics Society, Washington, USA, October 21-23, 2018, pp. 2516–2521, DOI: 10.1109/IECON.2018.8591399.

[25] Q. Li, V. Naing, J. A. Hoffer, D. J. Weber, A. D. Kuo, and J. M. Donelan, “Biomechanical energy harvesting: Apparatus and method,” in Proceedings of the IEEE International Conference on Robotics and Automation (ICRA), Pasadena, USA, May 19-23, 2008, pp. 3672–3677, DOI: 10.1109/ROBOT.2008.4543774.

[26] Q. Li, V. Naing, and J. M. Donelan, “Development of a biomechanical energy harvester,” Journal of NeuroEngineering and Rehabilitation, vol. 6, no. 22, 2009, DOI: 10.1186/1743-0003-6-22.

[27] R. Rarick, H. Richter, A. Van Den Bogert, D. Simon, H. Warner, and T. Barto, “Optimal design of a transfemoral prosthesis with energy storage and regeneration,” in Proceedings of the American Control Conference (ACC), Portland, USA, June 4-6, 2014, pp. 4108–4113, DOI: 10.1109/ACC.2014.6859051.

[28] F. Rohani, H. Richter, and A. J. Van Den Bogert, “Optimal design and control of an electromechanical transfemoral prosthesis with energy regeneration,” PLoS ONE, vol. 12, no. 11, pp. e0188266, 2017, DOI: 10.1371/journal.pone.0188266.

[29] M. R. Tucker and K. B. Fite, “Mechanical damping with electrical regeneration for a powered transfemoral prosthesis,” in Proceedings of the IEEE/ASME International Conference on Advanced Intelligent Mechatronics, Montreal, Canada, July 6-9, 2010, pp. 13–18, DOI: 10.1109/AIM.2010.5695828.

[30] H. Warner, D. Simon, and H. Richter, “Design optimization and control of a crank-slider actuator for a lower-limb prosthesis with energy regeneration,” in Proceedings of the IEEE International Conference on Advanced Intelligent Mechatronics, Banff, Canada, July 12-15, 2016, pp. 1430–1435, DOI: 10.1109/AIM.2016.7576971.

[31] E. Bolivar, S. Rezazadeh, and R. D. Gregg, “Minimizing energy consumption and peak power of series elastic actuators: A convex optimization framework for elastic element design,” IEEE/ASME Transactions on Mechatronics, vol. 24, no. 3, pp. 1334–1345, 2019, DOI: 10.1109/TMECH.2019.2906887.

[32] T. Elery, S. Rezazadeh, C. Nesler, J. Doan, H. Zhu, and R. D. Gregg, “Design and benchtop validation of a powered knee-ankle prosthesis with high-torque, low-impedance actuators,” in Proceedings of the IEEE International Conference on Robotics and Automation (ICRA), Brisbane, QLD, Australia, May 21-25, 2018, DOI: 10.1109/ICRA.2018.8461259.

[33] T. Elery, S. Rezazadeh, C. Nesler, and R. Gregg, “Design and validation of a powered knee-ankle prosthesis with high-torque, low-impedance actuators,” IEEE Transactions on Robotics, 2020.

[34] G. Lv, H. Zhu, and R. D. Gregg, “On the design and control of highly backdrivable lower-limb exoskeletons: A discussion of past and ongoing work,” IEEE Control Systems Magazine, vol. 38, no. 6, pp. 88–113, 2018, DOI: 10.1109/MCS.2018.2866605.

[35] H. Zhu, J. Doan, C. Stence, G. Lv, T. Elery, and R. Gregg, “Design and validation of a torque dense, highly backdrivable powered knee-ankle orthosis,” in Proceedings of the IEEE International Conference on Robotics and Automation (ICRA), Singapore, May 29 - June 3, 2017, DOI: 10.1109/ICRA.2017.7989063.

[36] H. Zhu, C. Nesler, N. Divekar, M. Taha Ahmad, and R. D. Gregg, “Design and validation of a partial-assist knee orthosis with compact, backdrivable actuation,” in Proceedings of the IEEE International Conference on Rehabilitation Robotics (ICORR), Toronto, ON, Canada, June 24-28, 2019, DOI: 10.1109/ICORR.2019.8779479.

[37] G. Khademi, H. Richter, and D. Simon, “Multi-objective optimization of tracking/impedance control for a prosthetic leg with energy regeneration,” in Proceedings of the IEEE Conference on Decision and Control, Las Vegas, USA, December 12-14, 2016, pp. 5322–5327, DOI: 10.1109/CDC.2016.7799085.

[38] G. Khademi, H. Mohammadi, H. Richter, and D. Simon, “Optimal mixed tracking/impedance control with application to transfemoral prostheses with energy regeneration,” IEEE Transactions on Biomedical Engineering, vol. 65, no. 4, pp. 894–910, 2018, DOI: 10.1109/TBME.2017.2725740.

[39] R. Riemer and A. Shapiro, “Biomechanical energy harvesting from human motion: Theory, state of the art, design guidelines, and future directions,” Journal of NeuroEngineering and Rehabilitation, pp. 22, 2011, DOI: 10.1186/1743-0003-8-22.

[40] A. M. Simon, N. P. Fey, K. A. Ingraham, S. B. Finucane, E. G. Halsne, and L. J. Hargrove, “Improved weight-bearing symmetry for transfemoral amputees during standing up and sitting down with a powered knee-ankle prosthesis,” Archives of Physical Medicine and Rehabilitation, vol. 97, no. 7, pp. 1100–1106, 2016, DOI: 10.1016/j.apmr.2015.11.006.

[41] F. Gao, F. Zhang, and H. Huang, “Investigation of sit-to-stand and stand-to-sit in an above knee amputee,” in Proceedings of the Annual International Conference of the IEEE Engineering in Medicine and Biology Society (EMBC), Boston, USA, August 30 - September 3, 2011, pp. 7340–7343, DOI: 10.1109/IEMBS.2011.6091712.

[42] M. K. Shepherd and E. J. Rouse, “Design and characterization of a torque-controllable actuator for knee assistance during sit-to-stand,” in Proceedings of the Annual International Conference of the IEEE Engineering in Medicine and Biology Society (EMBC), Orlando, USA, August 16-20, 2016, pp. 2228–2231, DOI: 10.1109/EMBC.2016.7591172.

[43] M. K. Shepherd and E. J. Rouse, “Design and validation of a torque-controllable knee exoskeleton for sit-to-stand assistance,” IEEE/ASME Transactions on Mechatronics, vol. 22, no. 4, pp. 1695–1704, 2017, DOI: 10.1109/TMECH.2017.2704521.

[44] J. Vantilt, K. Tanghe, M. Afschrift, A. Bruijnes, K. Junius, J. Geeroms, E. Aertbeliën, F. De Groote, D. Lefeber, I. Jonkers, and J. De Schutter, “Model-based control for exoskeletons with series elastic actuators evaluated on sit-to-stand movements,” Journal of NeuroEngineering and Rehabilitation, vo. 16, pp. 65, 2019, DOI: 10.1186/s12984-019-0526-8.

[45] H. A. Varol, F. Sup, and M. Goldfarb, “Powered sit-to-stand and assistive stand-to-sit framework for a powered transfemoral prosthesis,” in Proceedings of the IEEE International Conference on Rehabilitation Robotics (ICORR), Kyoto, Japan, June 23-26, 2009, pp. 645–651, DOI: 10.1109/ICORR.2009.5209582.

[46] M. Wu, M. R. Haque, and X. Shen, “Sit-to-stand control of powered knee prostheses,” in Proceedings of the ASME Design of Medical Devices Conference, Minneapolis, USA, 2017, pp. V001T05A015, DOI: 10.1115/DMD2017-3507.

[47] V. Norman-Gerum and J. McPhee, “Constrained dynamic optimization of sit-to-stand motion driven by Bézier curves,” ASME Journal of Biomechanical Engineering, vol. 140, no. 12, pp. 121011, 2018, DOI: 10.1115/1.4041527.

[48] R. S. Razavian, S. Greenberg, and J. McPhee, “Biomechanics Imaging and Analysis,” Encyclopedia of Biomedical Engineering, pp. 488–500, 2019, DOI: 10.1016/B978-0-12-801238-3.99961-6.

[49] B. Laschowski and J. McPhee, “Body segment parameters of Paralympic athletes from dual-energy X-ray absorptiometry,” Sports Engineering, vol. 19, no. 3, pp. 155–162, 2016, DOI: 10.1007/s12283-016-0200-3.

[50] B. Laschowski and J. McPhee, “Quantifying body segment parameters using dual-energy X-ray absorptiometry: A Paralympic wheelchair curler case report,” Procedia Engineering, vol. 147, pp. 163–167, 2016, DOI: 10.1016/j.proeng.2016.06.207.

[51] P. De Leva, “Adjustments to Zatsiorsky-Seluyanov’s segment inertia parameters,” Journal of Biomechanics, vol. 29, no. 9, pp. 1223–1230, 1996, DOI: 10.1016/0021-9290(95)00178-6.

[52] S. Seok, A. Wang, M. Y. Chuah, D. Otten, J. Lang, and S. Kim, “Design principles for highly efficient quadrupeds and implementation on the MIT cheetah robot,” in Proceedings of the IEEE International Conference on Robotics and Automation, Karlsruhe, Germany, May 6-10, 2013, DOI: 10.1109/ICRA.2013.6631038.

[53] S. Seok, A. Wang, M. Y. Chuah, D. J. Hyun, J. Lee, D. M. Otten, J. H. Lang, and S. Kim, “Design principles for energy-efficient legged locomotion and implementation on the MIT cheetah robot,” IEEE/ASME Transactions on Mechatronics, vol. 20, no. 3, pp. 1117–1129, 2015, DOI: 10.1109/TMECH.2014.2339013.

